# Unusual physiological properties of two ganglion cell types in primate retina

**DOI:** 10.1101/496455

**Authors:** Colleen E. Rhoades, Nishal P. Shah, Michael B. Manookin, Nora Brackbill, Alexandra Kling, Georges Goetz, Alexander Sher, Alan M. Litke, E. J. Chichilnisky

**Affiliations:** Department of Bioengineering, Stanford University, Stanford, CA 94305, USA; Department of Electrical Engineering, Stanford University, Stanford, CA 94305, USA; Department of Ophthalmology, University of Washington, Seattle, WA 98195, USA; Department of Physics, Stanford University, Stanford, CA 94305, USA; Department of Neurosurgery, Stanford University, Stanford, CA 94305, USA; Santa Cruz Institute for Particle Physics, University of California Santa Cruz, Santa Cruz, CA 95064, USA; Department of Ophthalmology Stanford University, Stanford, CA 94305, USA; Hansen Experimental Physics Laboratory, Stanford University, Stanford, CA 94305, USA

**Author notes:** Lead contact: Colleen E. Rhoades (Correspondence).

## Abstract

The visual functions of the diverse retinal ganglion cell types in the primate retina, and the parallel visual pathways they initiate, remain poorly understood. Here, the unusual physiological and computational properties of the ON and OFF smooth monostratified (SM) ganglion cells are explored. Large-scale multi-electrode recordings from 48 macaque retinas revealed that these cells exhibited strikingly irregular receptive field structure composed of spatially segregated hotspots, quite different from the receptive fields of previously described retinal ganglion cell types. The ON and OFF SM cells are paired cell types, but OFF SM cells exhibited stronger hotspot structure than ON cells. Targeted visual stimulation and computational inference demonstrate strong nonlinear subunit properties of each hotspot that contributed to the signaling properties of SM cells. Analysis of shared inputs to neighboring SM cells indicated that each hotspot could not be explained by an individual presynaptic input. Surprisingly, visual stimulation of different hotspots produced subtly different spatiotemporal spike waveforms in the same SM cell, consistent with a dendritic contribution to hotspot structure. These findings point to a previously unreported nonlinear mechanism in the output of the primate retina that contributes to signaling spatial information.

## Introduction

In the mammalian retina, a diverse collection of retinal ganglion cell (RGC) types extracts features of the visual scene and transmits the results to various targets in the brain. Each RGC type exhibits characteristic light responses, connects to specific retinal interneuron types, and covers the entire visual field, forming a distinct channel of information. Work in mice and other species has begun to reveal the diverse computations performed by the various RGC types, and their relationship to visual behaviors (Gollisch and Meister, 2010; Masland, 2001; Rodieck, 1998; Wässle, 2004).

However, in the primate retina, despite the existence of a nearly complete anatomical catalog of roughly 20 RGC types, the understanding of their distinct visual computations and underlying cellular and circuit properties remains limited (Dacey et al., 2003; Yamada et al., 2005). Most physiological studies have been performed on the five numerically dominant primate RGC types, ON and OFF midget (Dacey, 1993a), ON and OFF parasol (Chichilnisky and Kalmar, 2002), and small bistratified (Chichilnisky and Baylor, 1999; Dacey, 1993b; Field et al., 2007), which make up about 75% of the visual signal. These cells are usually characterized as exhibiting classical Gaussian center-surround receptive field (RF) structure, with relatively little evidence for specialized functional properties such as those found in mouse RGCs (Crook et al., 2008; Enroth-Cugell and Robson, 1966; Kuffler, 1953; Rodieck, 1998), but see (Manookin et al., 2018). The function of visual signaling in the remaining low-density RGC types remains largely unknown (Puller et al., 2015). Notably, the most specific computations found in mouse RGCs (e.g. direction selectivity) have not yet been observed in primates (Barlow and Levick, 1965; Barlow et al., 1964; Puller et al., 2015), and the anatomical homology of RGC types between rodents and primates is far from clear (Roska and Meister 2014). Thus, it is uncertain whether the poorly understood RGC types in primates could serve distinctive roles in vision based on unique physiological mechanisms, as is the case in other species.

A primary reason for this limited understanding is the technical challenge of recording from low-density RGC types in primate, each of which constitute only a few percent of the total population (Dacey et al., 2003; Yamada et al., 2005). This study combines single-cell physiological recording with large-scale multi-electrode recording from hundreds of RGCs in isolated primate retina to explore the responses of two low-density RGC types—the ON and OFF smooth monostratified (SM) cells. Quantitative analysis of SM cell light responses, relative to those of simultaneously-recorded parasol cells, enabled their identification across recordings from different retinas. Both SM cell types exhibited irregular RFs with multiple distinct hotspots of light sensitivity, which are not observed in the high-density cell types. Analysis of neighboring SM cells suggested that the hotspots could not be simply explained by a single shared presynaptic input. An unexpected spike generation mechanism was revealed by visual stimulation of individual hotspots producing distinct spike waveforms in a given RGC. Closed-loop visual stimulation and computational inference identified that the hotspots behave as nonlinear subunits that are more distinct and spatially segregated than those found in other cell types and species.

## Results

The results below show that (1) SM cells can be reliably recorded and identified, (2) their RFs exhibit an unusual structure composed of distinct hotspots, (3) the hotspots are associated with different spike waveforms, and (4) a computational model that extracts the hotspots as nonlinear subunits is predictive of the RGC responses.

### Receptive field properties of simultaneously recorded retinal ganglion cell types

To explore the properties of poorly understood RGC types, large-scale multi-electrode recordings were used to simultaneously record the light responses of hundreds of RGCs (Chichilnisky and Kalmar, 2002; Field et al., 2010; Frechette et al., 2005; Litke et al., 2004). The spatial, temporal, and chromatic response properties of each recorded RGC were examined by computing the reverse correlation between its spike train and a spatiotemporal noise stimulus. The resulting spike-triggered average (STA) stimulus captures the spatial RF, time course, and chromatic properties of each cell analyzed (Fig. 1; Chichilnisky, 2001). To distinguish cell types, the RF area and first principal component of the time course were examined (Fig. 1A). As previously demonstrated (Field et al., 2007), the five major high-density cell types (ON parasol, OFF parasol, ON midget, OFF midget, and small bistratified) can be readily identified based on these properties (Fig. 1D-H). In addition, in a given recording, a collection of RGCs was identified that had RF sizes and other functional properties clearly distinct from the five major types (Fig. 1B-C). Often, one or more subsets of these RGCs appeared to comprise a single functional type, based on their homogeneous RF size, response time course, and spike train autocorrelation properties, as well as non-overlapping RFs, consistent with the mosaic organization known for most RGC types (Chichilnisky and Kalmar, 2002; Dacey, 1993a; Devries and Baylor, 1997; Field et al., 2007; Frechette et al., 2005; Peichl, 1991; Wassle et al., 1981). Below, evidence is provided that two of the low-density RGC types frequently observed were ON and OFF SM cells.

**Figure 1:**
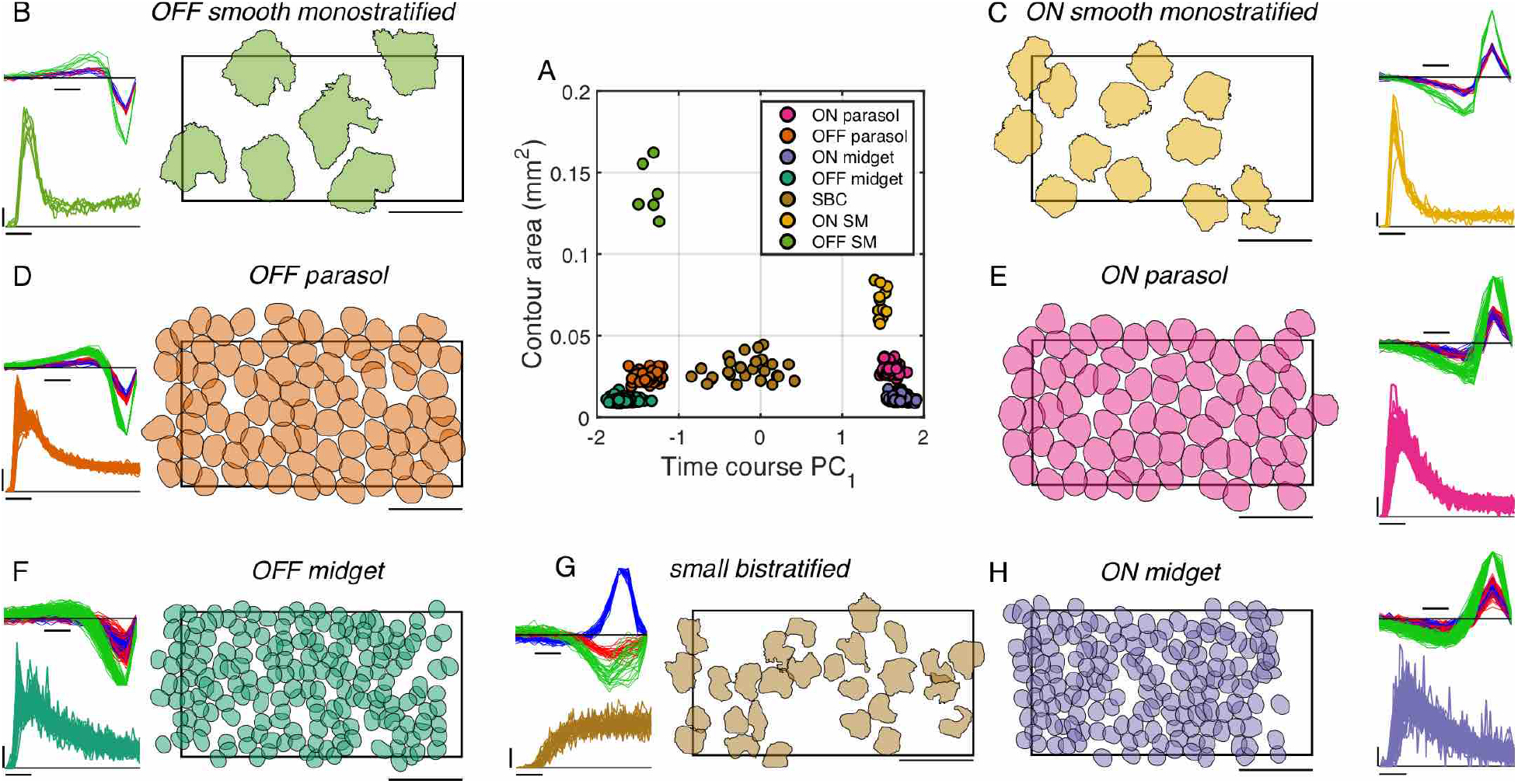
Receptive field properties of simultaneously recorded retinal ganglion cell types. A: The contour area and first principal component of the time course for each RGC in a single recording reveals distinct clusters, which are classified into seven unique cell types. Temporal equivalent eccentricity = 7 mm or 31.4 degrees (Dacey and Petersen, 1992; Perry and Cowey, 1985) B: Properties of a single class, anatomically identified as OFF smooth monostratified (see Fig. 2). Right: The contours outline the normalized RF at a threshold equal to 0.18 and then shrunk around the center of the RF (see Methods). The contours should not be interpreted as the actual RF sizes, but rather the relative sizes of the various RGC types. Rectangle indicates the outline of the multi-electrode array (1800 μm × 900 μm). Scale bar = 500 μm. Top left: The STA time courses for red, green, and blue display phosphors, normalized to align the peak amplitude for each cell. Scale bar = 50 ms. Bottom left: Autocorrelation function where the y axis indicates the probability of spiking (scale bar = 0.01), normalized to unit area, and the x axis represents time (scale bar = 10 ms). C-H: Properties of additional cell classes recorded simultaneously as described for panel B.

### Classification of low-density cell types across recordings

Identification of SM cells required a different logic from that used to identify the five major RGC types in previous studies. The high-density RGC types can be easily classified across recordings because of their unique density and spatial, temporal, and chromatic light response properties (Chichilnisky and Kalmar, 2002; Dacey, 1993a; Field et al., 2007). However, many of the anatomically identified low-density RGC types have similar dendritic field size and density to one another, and other physiological properties of these cell types are largely unknown, making classification difficult (Dacey et al., 2003). Thus, to identify low-density cell types across recordings, a method of compensating for variability across recordings was necessary.

Analysis was performed on 53 recordings from 40 monkeys, each of which contained multiple ON and/or OFF low-density cell types. First, within each recording, the collection of low-density RGC types was segregated into several distinct groups of cells, based on differences in their time courses and autocorrelation function (as described in Fig. 1). Second, separately for the ON and OFF low-density RGCs in each recording, the group of cells with the fastest time course, as measured by the time of zero crossing, was labeled as α, while the remaining low-density cells were lumped together and labeled β (Fig. 2A, D). Third, the times of zero crossing and biphasic index values (Fig. 2A) of all the low density cells were normalized by those of the simultaneously recorded ON and OFF parasol cells, respectively. After these steps, the α group of cells, across many recordings, had homogeneous times of zero crossing and biphasic index values: the average distance from an ON (OFF) α cell to the center of the α cluster is 0.13 (0.21) (Fig. 2B, C). The ON (OFF) midget RGCs, analyzed with the same method, showed a much higher average distance of 0.51 (0.78). The tight clustering of α suggested that it represented a single cell type identified across recordings. A support vector machine classifier was used to conservatively define a boundary around the α cells. The β cluster exhibited greater variability, and based on observed RF overlap, probably consisted of multiple RGC types. β cells will not be analyzed further.

**Figure 2:**
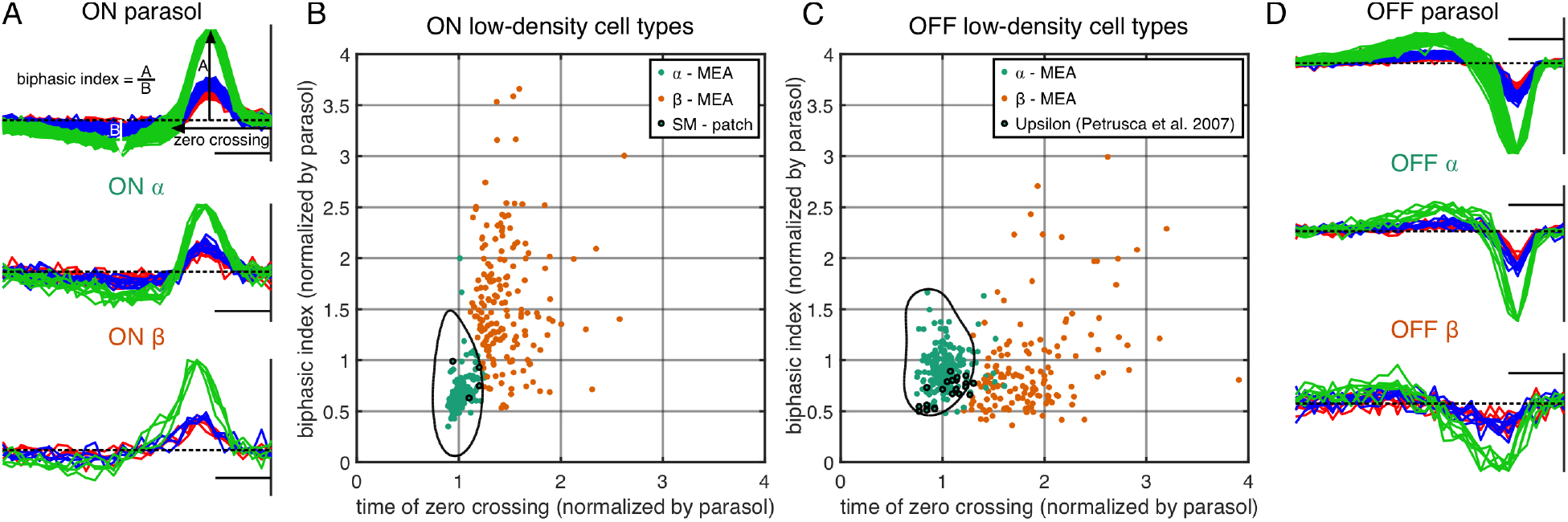
Classification of low-density cell types across recordings. A: The time courses, normalized by the maximum amplitude, of a population of simultaneously recorded ON parasol cells and ON low-density cell types. The measurement of zero crossing and biphasic index are indicated. The vertical line at the end of the time course indicates the time of the spike. Scale bar = 50 ms. B: Each point indicates the time of zero crossing and biphasic index of an ON low-density cell divided by the average respective properties of the ON parasol cells in each recording. The fastest low-density cell type in each recording (α; teal) cluster together across the 384 cells from 31 ON cell recordings while the remaining low-density cell types (β; orange) recorded in the same preparations are distinct. Within the black boundary, a support vector machine predicts a classification of α with greater than 95% probability. Only the cells within the black boundary will be included in further analysis. The outlined black dots correspond to anatomically identified ON SM cells, which all lie within the black boundary. C: As described in B but for 346 cells from 27 recordings containing multiple types of OFF low-density cells. The 23 black outlined cells correspond to upsilon cells recorded on an MEA and reported in Petrusca et al., 2007. D: The normalized time courses of populations of simultaneously recorded OFF parasol cells and two OFF low-density cell types as described in A.

The ON and OFF α clusters were hypothesized to correspond to the ON and OFF smooth monostratifed (SM) cells, for several reasons. First, the ON and OFF α cells had very similar properties to one another, other than response polarity, suggesting that they correspond to paired cell types (Ravi et al., 2018; Watanabe and Rodieck, 1989). Among the known low-density primate RGC types, other than ipRGCs and cells that receive strong input from S cones (Dacey, 2004; Dacey et al., 2005; Liao et al., 2016), only two are unambiguously paired: SM and narrow thorny. Second, the response kinetics of the ON and OFF α cells were very similar to those of parasol cells, with normalized biphasic index and zero crossing values close to 1 (Fig. 2B, C). SM cells, unlike narrow thorny cells, co-stratify with the parasol cells in the inner plexiform layer, in a region known for more transient kinetic response properties (Crook et al., 2008). Third, the RF sizes of α cells match the previously reported RF sizes of SM cells – approximately 2-3 times that of parasol cells (Crook et al., 2008; Petrusca et al., 2007). For these reasons, ON and OFF α cells were tentatively identified as ON and OFF SM cells, respectively.

To confirm this hypothesized anatomical identity, the same time course measurements were performed on morphologically identified ON SM RGCs and ON parasol RGCs with single-cell patch clamp recordings. Four ON SM cells from three retinas showed similar properties to α cells recorded on the MEA (Fig. 2B: black outlined cells).

Note that the response properties of 21 out of 23 of previously reported “upsilon” cells, hypothesized to be SM cells (Petrusca et al., 2007), fall within the classification boundary (Fig. 2C: black outlined cells).

### Receptive field structure of ON and OFF smooth monostratified cells

SM cells displayed surprising, patchy RF structure not observed in other simultaneously recorded RGC types. The RF structure was examined using coarse spatiotemporal noise stimuli because finer stimuli elicited weak responses in SM cells. To avoid lattice artifacts introduced by large stimulus pixels (84.8 × 84.8 um), the stimulus was jittered on a finer underlying lattice (5.3 × 5.3 um). The resulting RFs were irregularly shaped, with large distinct hotspots separated by regions of low sensitivity (Fig. 3B). Analyzed at a comparable spatial scale (~12 pixels covering the RF), simultaneously recorded parasol cells exhibited RF structure well approximated by a Gaussian form (Chichilnisky and Kalmar, 2002). Similar hotspots were observed in anatomically identified ON SM cells recorded with patch clamp recording (Fig. 3B; bottom left). The unusual SM cell RF structure was quantified by an inhomogeneity index, given by the mean squared deviation from a 2D Gaussian fit to the RF (after normalizing, masking out the background with a threshold of 0.3, and correcting for variance in the STA due to measurement noise). In 97 SM cells from six recordings and 830 parasol cells from three recordings, the SM cell RFs exhibited an inhomogeneity index (0.047 ± 0.028; mean ± SD) significantly higher than that of parasol cell RFs (0.012 ± 0.011) (Wilcoxon rank sum test p < 1e-10). The hotspot structure in OFF SM cells (0.065 ± 0.027) was also more pronounced than that in ON SM cells (0.033 ± 0.020), resulting in a higher inhomogeneity index and consistent with visual inspection (see Fig. 3B; Wilcoxon rank sum test p< 1e-7 in 55 ON cells and 42 OFF cells from 6 recordings).

**Figure 3:**
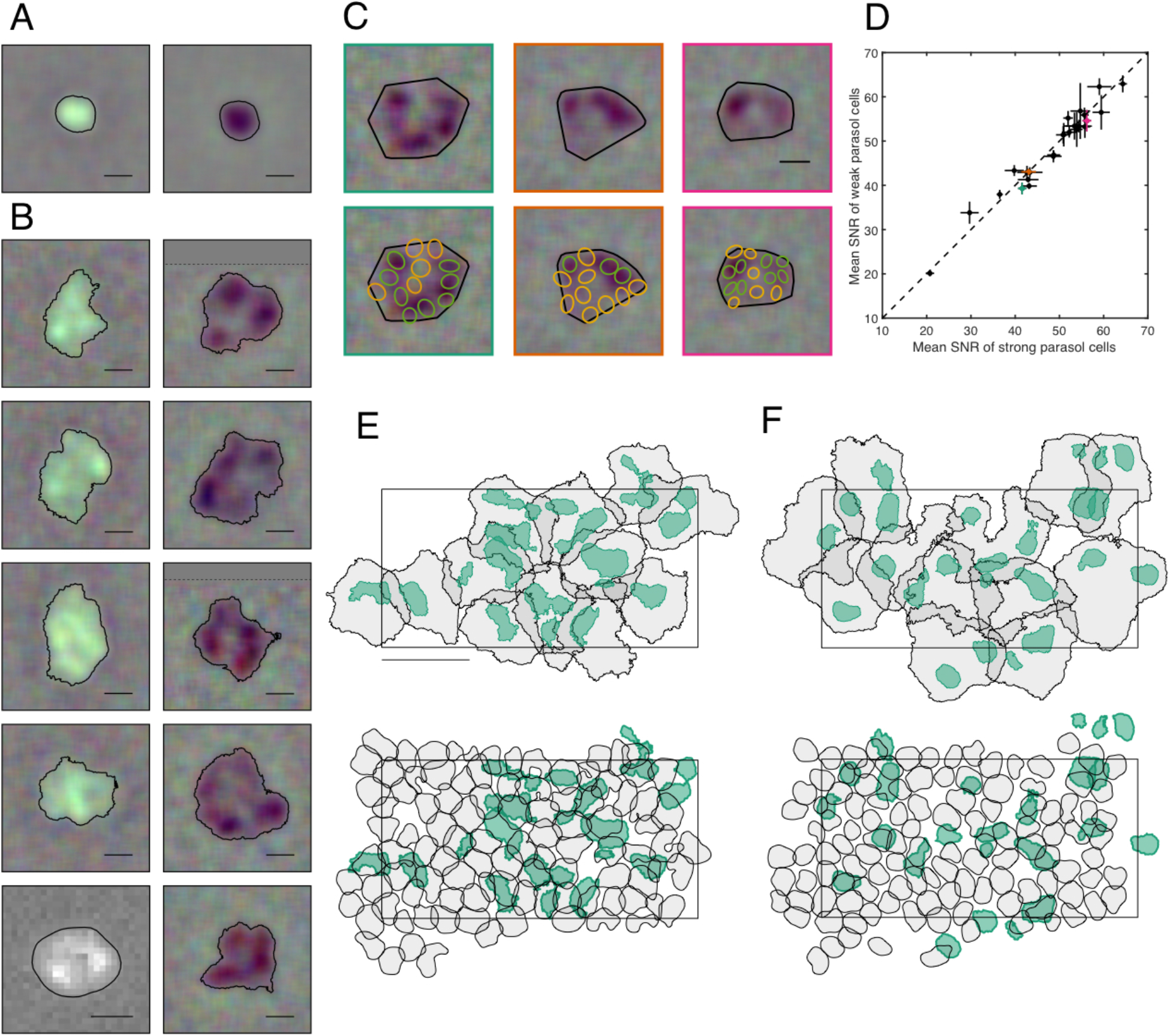
Receptive field structure of ON and OFF smooth monostratifed cells. A: ON (left) and OFF (right) parasol cells do not exhibit hotspots in their RFs. Contour threshold = 0.15. Scale bar = 200 μm. B: The ON (left column) and OFF (right column) SM cells have RFs with distinct hotspots. The bottom left ON SM cell was recorded with patch clamp while the remaining were recorded on an MEA. The dotted line indicates the end of the visual stimulus and the remainder of the STA was filled in with grey. Contour threshold = 0.15. Scale bar = 200 μm. C: RF alignment of three OFF SM cells (top) with the simultaneously recorded OFF parasol cells (bottom; green/yellow). The green parasol cells cover the hotspots in the SM RF while the yellow parasol cells fall between the hotspots. Scale bar = 200 μm. D: The SNR of the simultaneously measured parasol cells over the SM hotspots (green parasol cells) and in between the hotspots (yellow parasol cells) was compared: mean of all strong/weak parasol cells ± SEM for 26 SM cells from three recordings. Three examples are highlighted in teal, orange, and pink and correspond to the three SM cells shown in C. E: The RFs of a population of ON SM cells (black; contour threshold = 0.1) and the hotspot(s) within each RF (teal; variable threshold). Below the same hotspots are shown (teal) with the simultaneously recorded ON parasol cells (contour threshold = 0.5). The rectangle indicates the position of the electrode array. Temporal equivalent eccentricity = 12 mm. Scale bar = 500 μm. F: As described in E for OFF SM and OFF parasol cells.

The low light sensitivity in the gaps between RF hotspots could not be explained by damaged photoreceptors. This was determined by examination of simultaneously recorded parasol RGCs with RFs overlying the RFs of SM cells. Specifically, for each SM cell, the OFF parasol cells with RFs intersecting the SM cell RF (determined by the convex hull of a contour; see Methods) were identified. The parasol cells were then divided into two groups, according to whether the parasol cell RF overlaid a strong region in the SM cell RF (i.e. hotspots) or weak region (i.e. between hotspots). If the SM cell hotspots were the result of photoreceptor damage, then the parasol cell RFs overlying hotspots (Fig. 3C; green) would be expected to have higher signal to noise ratio (SNR) than the parasol cell RFs overlying gaps between hotspots (Fig. 3C; yellow) because the same photoreceptors provide input to both cell types. Contrary to this prediction, no systematic difference was observed in the SNR of the parasol cells over the strong and weak areas of the SM cell RF (Fig. 3D).

Another possibility, given the size of the hotspots, is that the SM cells are in fact collections of parasol cells erroneously grouped together. Simultaneous recording of SM cells and parasol cells enabled direct comparison of the hotspot locations to the parasol cell RFs. The contour threshold for each SM cell was adjusted so that the average contour area of the hotspots approximated the average area of the parasol cells. While some hotspots overlapped with parasol cells (as would be expected by chance), the SM cell hotspots are not composed of parasol cell RFs (Fig. 3E-F). Moreover, the same hotspot structure was observed in patch clamp recordings from single, morphologically verified SM cells (Fig 3B bottom left), indicating that this unique property is not a result of recording artifacts.

### Coordination of fine RF structure in neighboring RGCs

Fine-scale interdigitation of RFs, resulting in complete tiling of visual space more uniformly than would be expected from a lattice of independently formed RFs, has been observed in other RGC types (Gauthier et al., 2009). To test if ON and OFF SM cell RFs interdigitate similarly, the uniformity index (UI; Gauthier et al., 2009), computing how uniformly each recorded population of SM cells covered the visual field, was analyzed. The UI was given by the fraction of pixels within the convex hull of the mosaic that were covered by the RF contour of exactly one cell. The contour level for the normalized RFs was set to maximize the UI, thus balancing the area covered by no cells with the area covered by multiple cells. To test whether the observed interdigitation exceeded chance expectations, the UI of the data (Fig. 4A) was compared to a null distribution of UIs (Fig. 4C) resulting from rotating each cell randomly around the center of its contour (Fig. 4B). The UI of the data was in general at the extreme high end of the distribution obtained from random rotations; e.g. above the 98th percentile in 9/10 preparations (Fig. 4D), confirming that the ON and OFF SM cell types interdigitated.

**Figure 4:**
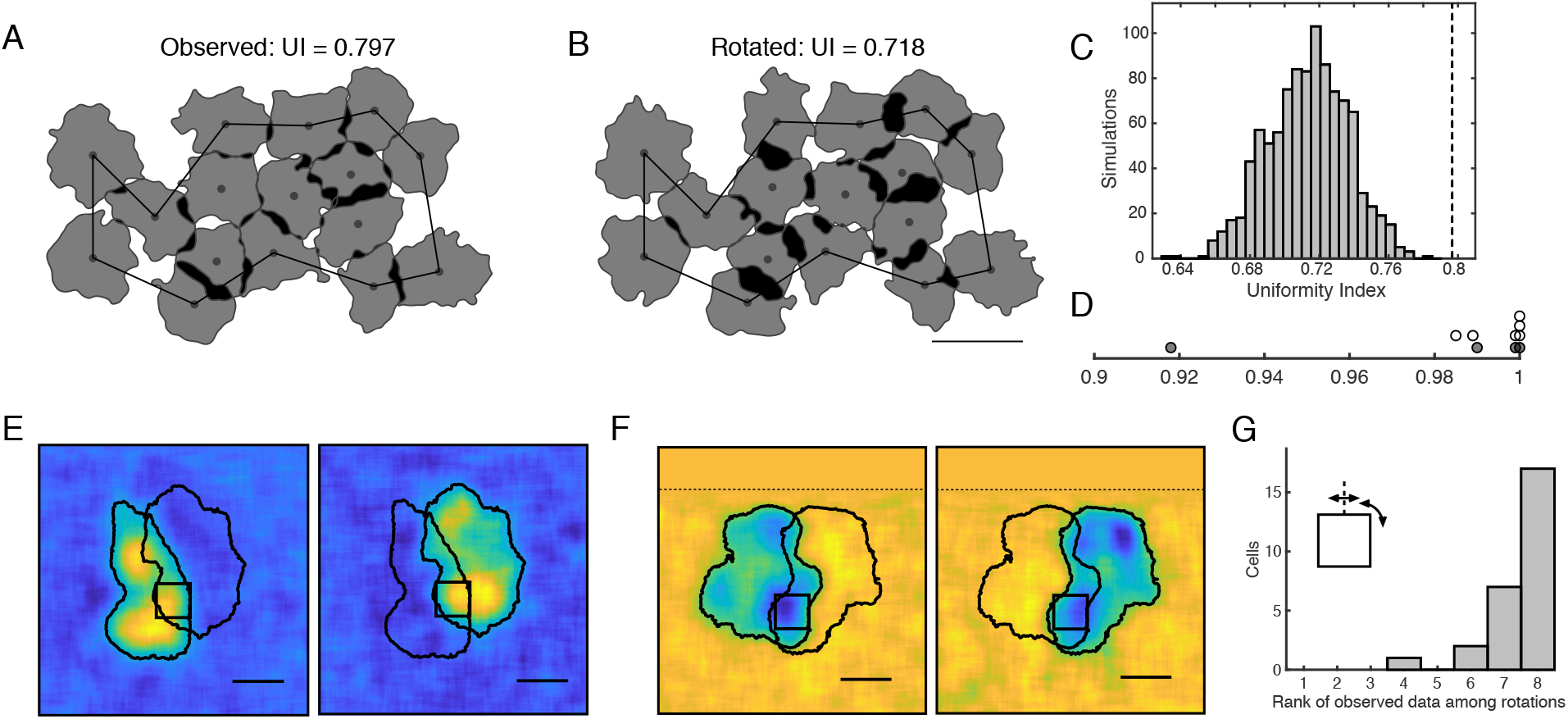
Coordination of fine RF structure in neighboring RGCs. A: Contours outlining the RFs of a population of SM cells showed interdigitation of neighboring cells. White regions are covered by no cells, gray by one cell, and black by multiple cells. The Ul was computed within the black boundary (see Methods). Contour threshold = 0.13. B: Each cell was rotated at a random angle around its center point and the contour threshold was recomputed to maximize the UI. Scale bar = 500 μm. C: The UI of the observed data (black dotted line) was compared to a null distribution composed of 1000 samples with randomly rotated cells. D: Across 6 ON SM mosaics (white circles) and 4 OFF SM mosaics (grey circles), the percentile of the UI of the observed data compared to 1000 trials of randomly rotated data is reported. E: Two ON SM cells that partially share a hotspot, are shown with the RFs at contour threshold = 0.2. The black square shows the 132.5 × 132.5 μm region used for rotation/reflection analysis. Scale bar = 200 μm. F: Same as E for two OFF SM cells. G: For 27 cell pairs, the rank of the correlation between the region of overlap for the observed data among the seven rotations and reflections showed the observed data skewed lower in correlation than the transformed data. If cell pairs completely shared hotspots the rank of the observed data would skew towards 1.

A possible mechanism for the irregular RF structure is that each hotspot represents the input of a single presynaptic neuron, such as a bipolar or amacrine cell. Two observations argue against an origin from individual bipolar cells. First, the hotspots are large compared to RFs expected from known primate bipolar types (Boycott and Wassle, 1991; Tsukamoto and Omi, 2015, 2016), but see (Dacey et al., 2000). Second, SM cells exhibited significant frequency-doubled responses to contrast-reversing gratings with spatial periods ~50-100 μm (Crook et al., 2008; Petrusca et al., 2007) smaller than or comparable to the diameter of the hotspots (~100 μm; see Fig. 3B). Thus, each hotspot is likely composed of multiple bipolar cells.

More generally, if the hotspots were produced by input from a single cell of any kind, then RFs of neighboring SM cells should either share entire hotspots, or not overlap at all. While the majority of SM cell RFs did not overlap at all (Fig. 3E, F), a small subset of SM cells exhibited significant overlap with a neighbor, enabling an analysis of whether the hotspot shape was the same in the two cells. 27 SM cell pairs from 15 recordings that had measurable overlap were examined. A 132.5 μm by 132.5 μm region (black square; Fig. 4E, F) was identified and centered on the pixel with the maximum intensity product in the RFs of the two cells – the peak of the shared input region. Under the assumption that the hotspots were produced by input from a single cell, the hotspot would be expected to have the same spatial structure in the RFs of both cells. Therefore the spatial similarity of the RFs within the region of overlap would be reduced by rotations and reflections (lower ranks; Fig. 4G). Instead, rotations and reflections around the midpoint of the region of overlap generally increased the correlation between the RF pixels of the two cells (higher ranks; Fig. 4G). This finding supports the idea that the hotspots were not formed by the input of individual presynaptic cells.

### Potential mechanism of RF hotspots

Another mechanism that could contribute to hotspots is active conductances or other electrical properties of SM cell dendrites. This possibility is suggested by a surprising observation: visual stimulation of each hotspot generated a slightly different spike waveform in each SM cell. These waveforms originate from spatially distinct regions, which correspond to the locations of the hotspots.

In the process of segregating spikes from different RGCs based on their spike waveforms (spike sorting – see Methods), several slightly different spike waveforms were often observed for a single SM cell (Fig. 5A, C). These distinct waveforms were confirmed with spectral clustering, indicating that several reliably different spike waveforms could be produced by a single SM cell. Moreover, when the STA was computed using reverse correlation with just the spikes in one sub-cluster, only one or two hotspots from the SM cell RF emerged (Fig. 5B).

**Figure 5:**
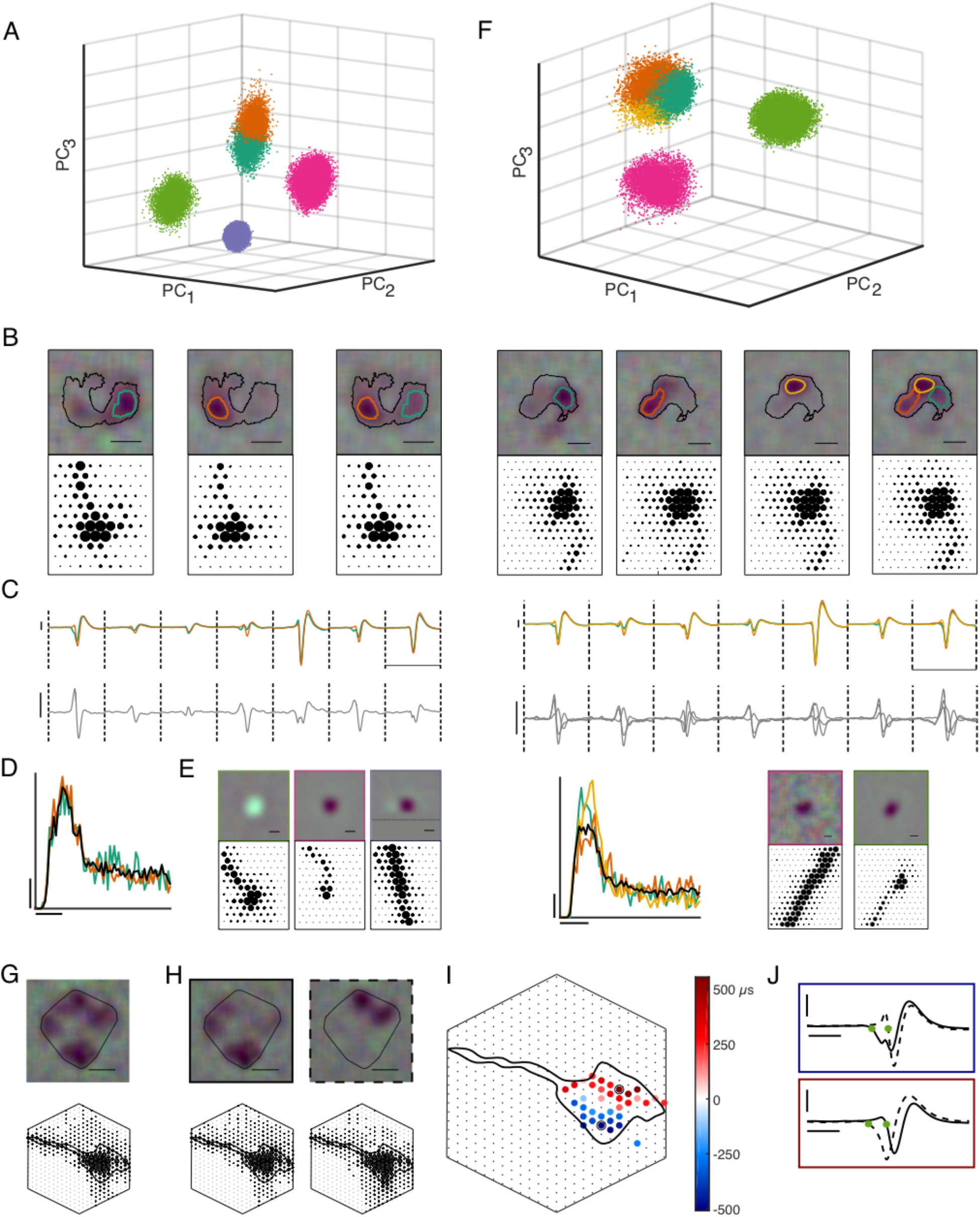
Potential mechanism of receptive field hotspots. A: The first three principal components of the spike waveforms from several cells recorded on a particular electrode. Each cluster corresponds to spikes from a different cell with the orange and teal cluster representing spikes from an OFF SM cell. Additional recorded waveforms that could not be reliably associated with a specific neuron, according to the inclusion criteria (see Methods) are omitted for clarity. B: The RF and EI computed from spikes in the teal and orange clusters followed by the combined cluster. The black contour is fit on the full SM cell and the teal and orange contours are fit at a threshold of 0.2. Scale bar = 200 μm. Electrode spacing = 60 μm. C: The mean waveforms in the SM cell sub-clusters across the seven electrodes used in spike sorting (top). The difference between the mean waveforms is shown on an expanded scale (bottom). The y axis is voltage difference with a scale bar of 50 μV and the x axis is time with a scale bar of 5 ms. D: The autocorrelation function of the SM cell (black), and of each SM cell sub-cluster (orange and teal). There is no refractory violation when the subcluster spikes are combined and each autocorrelation function has the same shape. The y axis indicates the probability of spiking (scale bar = 0.01), normalized to unit area, and the x axis represents time (scale bar = 10 ms). E: The RF and EI computed from spikes on non-SM cells recorded on this electrode, outlined in the color corresponding to the cluster in A. The electrical images do not match the SM cell. The dotted line indicates the end of the visual stimulus and the remainder of the STA was filled in with grey. Scale bar = 100 μm F: Same as A-E from a different recording with three subclusters (teal, orange, brown) of the SM cell. G: The RF and maximum projection of the EI of an OFF SM cell. The contour threshold was 0.25 and is shown as the convex hull. Scale bar = 200 μm. Electrode spacing = 30 μm. The hexagon indicates the outline of the 519-electrode multi-electrode array. H: The RF and maximum projection of the EI computed from clusters of spikes with different waveforms, which together make up the SM cell in G. Each waveform was associated with a different region of the RF, yet the EI remains the same. Scale bar = 200 μm. Electrode spacing = 30 μm. I: Each electrode was colored based on the timing difference between when electrical activity began in each cluster. The red electrodes indicates when signal is present for the second cluster (H, dotted border) before the first cluster (H, solid border) and vice versa for the blue electrodes. The RF produced by spikes generated by different waveforms (H) aligns with the observed origin of the EI (Also see movie S1). J: Two example electrodes with maximal timing differences, corresponding to the electrodes outlined in black in panel I. The blue (red) electrode example shows electrical propagation originating from the first (second) cluster. The timing differences reported in panel I were measured as the difference between the green asterisks. Horizontal scale bar = 1 ms. Vertical scale bar = 50 μV.

The concern that these sub-clusters could reflect spikes from multiple RGCs incorrectly identified as arising from a single SM cell was excluded for several reasons. First, if this were the case, then the collection of spikes in the larger cluster identified as an SM cell should exhibit refractory period violations, as is generally true when midget and parasol cell spikes are erroneously combined. However, the autocorrelation function of the resulting spike train (Fig. 5D) showed no spikes within 1.7 ms of each other. Second, the form of this autocorrelation was highly consistent across SM cells (Fig. 1), which would not occur if each SM cell cluster were composed of various erroneous mixtures of spikes from different cells of different types in the vicinity. Third, the electrical image (EI), or average spatial electrical footprint observed on the array during a spike (Greschner et al., 2014; Litke et al., 2004; Petrusca et al., 2007), was nearly identical for all sub-clusters (Fig. 5B), but was strikingly different for different RGCs (Fig. 5E). This is consistent with the fact that the EI reveals the locations of the soma, dendrites, and axons of the recorded cell (Litke et al., 2004). Fourth, given that the hotspots were roughly the size of parasol cell RFs (Fig. 3), it seems possible that the subclusters were in fact parasol cell spikes. However, the observed hotspots did not align to the RFs of simultaneously recorded parasol cells (Fig. 3E-F), and there are no other RGC types in the primate retina that occur at the density of parasol cells (Dacey, 2004). Finally, the possibility that SM cells represent mixtures of erroneously combined spikes from spiking amacrine cells was excluded both because their EIs are entirely different (Greschner et al., 2014; Litke et al., 2004; Petrusca et al., 2007), and because in cases when amacrine cells were recorded, the hotspots did not align with the amacrine cell RFs (not shown).

The multiple distinct spike waveforms observed in SM cells are not a ubiquitous feature of primate RGCs: waveform analysis of midget and parasol cells failed to reveal any sub-clusters of the kind seen in SM cells (not shown). The distinct spike waveforms were also not obviously attributable to inactivation of ion channels that would cause spikes closely spaced in time to have different waveforms: removal of all spikes within 500 ms of preceding spikes produced similar results (not shown). Thus, the distinct waveforms appear to represent an unusual feature of SM cells: the hotspots reflect its electrical spatial structure.

To investigate the relationship between the RF hotspots and the dendritic origin of the waveform differences, the EIs of SM cells were examined on a hexagonal array with 30 μm spacing (rather than 60 μm) for a higher resolution view of electrical propagation (Fig. 5G). To estimate the time of initial activity in the dendrites, the time when the waveform first fell below a threshold was measured for each clusters’ mean waveform on each electrode (Fig. 5J, green asterisk), Comparing this time for each cluster revealed spatial segregation in the appearance of the first electrical activity (Fig. 5I). For the cluster with hotspots in the bottom left (Fig 5H: solid outline), the electrical activity appeared first on the electrodes in the bottom left of the EI (Fig. 5I; blue electrodes), and for the cluster with hotspots in the top right (Fig 5H: dotted outline), the electrical activity appeared first on the electrodes in the top right of the EI (Fig. 5I; red electrodes). See Movie S1 to observe the electrical activity over time. Similar results were observed in 5 out of the 7 cells recorded with the high-density electrode arrays that could be used to test this effect.

### Nonlinearity of hotspots

What role do RF hotspots play in visual computation? One possibility is that they behave as nonlinear computational subunits, as has been observed with bipolar cell inputs to other RGCs (Demb et al., 1999, 2001; Hochstein and Shapley, 1976; Shapley and Victor, 1979; Victor and Shapley, 1979). To test for this possibility, a closed loop experiment was performed targeting the distinct spots independently. First, the RF of the SM cell was identified (Fig 6A). Then circular stimulus regions were placed over two hotspots, with their size adjusted to match the weight of the SM cell RF in each region. The two spots were modulated independently while measuring the firing rate of the SM cell, probing the interactions between hotspots. The temporally integrated contrast on each region (see Methods) and the firing rate of the SM cell were binned and fitted with a surface governed by either a linear model or a nonlinear model (Fig. 6C, E). In the linear model, visual inputs from the distinct regions are summed before a rectifying nonlinearity; in the nonlinear model, the input from each region is rectified before summation. For example, if one region was stimulated with a strong negative stimulus while the other region was stimulated with a strong positive stimulus, the linear model would predict a low firing rate, while the nonlinear model would predict a high firing rate. For each cell, the observed data (Fig. 6B) was compared to both fitted models. In 30 cells from six retinas, the nonlinear model resulted in lower mean squared deviation from the data, indicating that nonlinear mechanisms acting before the summation of signals across hotspots are important for explaining SM cell responses (Fig. 6E). Thus, hotspots behave as nonlinear computational subunits within the RF, potentially as a result of aggregating bipolar cell inputs (see below and Discussion).

**Figure 6:**
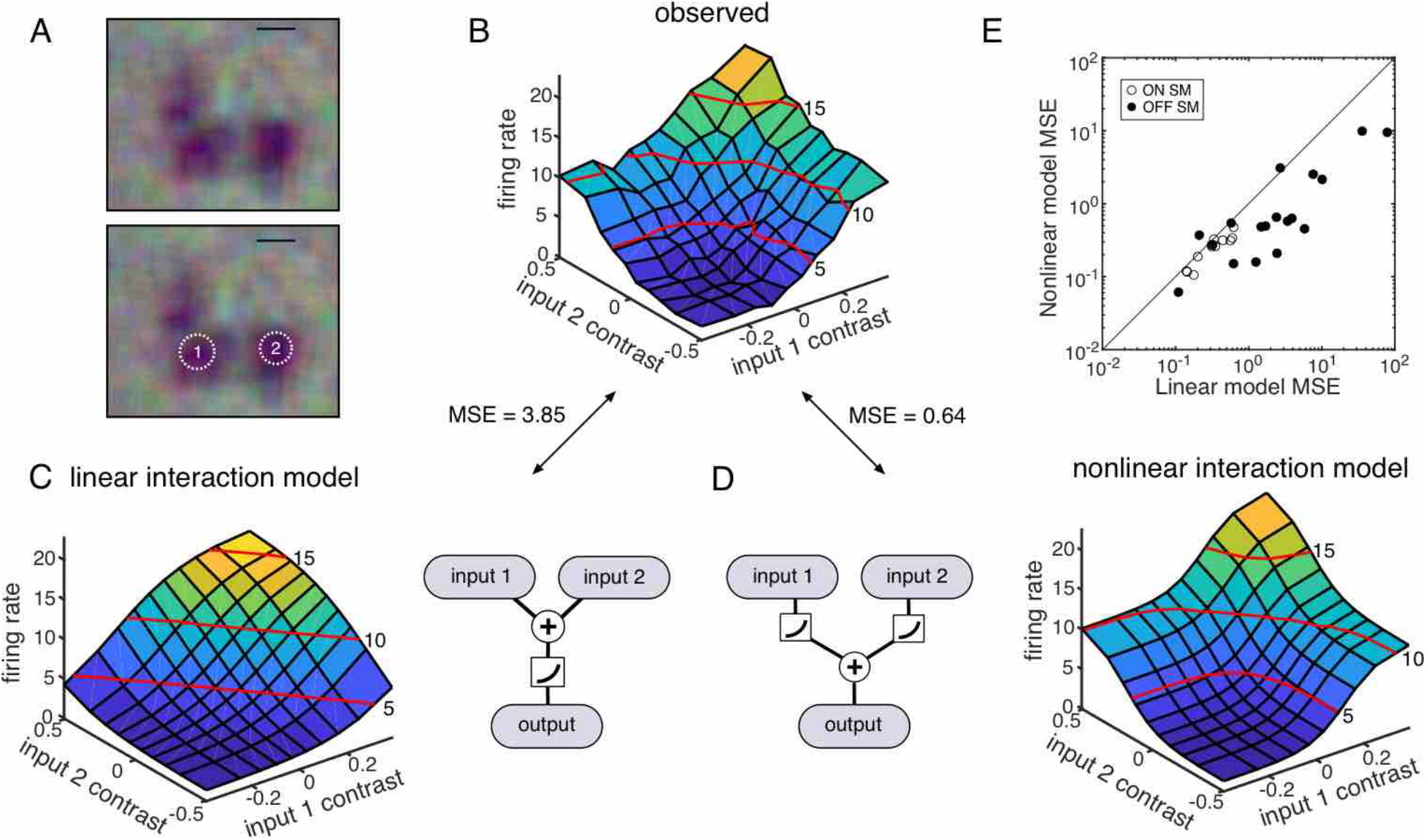
Nonlinearity of hotspots. A: The RF of an SM cell was computed with a white noise stimulus on an underlying jittered lattice. Two regions for targeted stimulation with equal weight in the RF are identified. Scale bar = 200 μm. B: The instantaneous contrast on each input region was computed and binned as a function of the firing rate of the SM cell. The red lines indicate the firing rate contour at 5, 10, and 15 Hz. The observed data more closely approximated the nonlinear model (D) with lower mean squared error. C: The predicted firing rate of a linear model as a function of the instantaneous contrast on each input region (left). The model was fit to the observed data, minimizing mean square error. A schematic of the linear model shows the input regions are summed before rectification (right). D: As in C, but for a nonlinear response model. The input regions were rectified and then summed (left). E: Across 30 cells and 6 retinas, the nonlinear model better approximated the combination of the input regions. Open circles correspond to ON SM cells and closed circles correspond to OFF SM cells.

### Subunit estimation through spike-triggered stimulus clustering

Given that the observed hotspots combine nonlinearly, a computational procedure that identifies the nonlinear subunits in the RF should also identify the RF hotspots (Liu et al., 2017). To test this prediction, a computational model designed to extract the spatial structure of nonlinear subunits was used (Shah et al., 2018).

The model consisted of two linear-nonlinear (LN) stages (Fig. 7A). In the first stage, the spatial stimulus was projected linearly onto a collection of spatial subunits, and the output of each subunit was subjected to an exponential nonlinearity. In the second stage, the inputs from the first stage were weighted and summed, and the output was subjected to a final nonlinearity. To reduce the dimensionality of the data, the subunits were assumed to be spatio-temporally separable and to each have the same temporal filter, estimated from the STA. The spatial stimulus associated with each spike was computed by convolving the SM cell response time course with the stimulus. Model fitting was performed on 80% of the data in two alternating stages: first the subunits were estimated by clustering the spike triggered stimuli, and second, the subunit weights and parameters of the output nonlinearity were estimated. This procedure yielded multiple estimated spatial subunits, determined by two hyperparameters: the number of subunits and a regularization value that emphasized spatial locality (see Methods).

**Figure 7:**
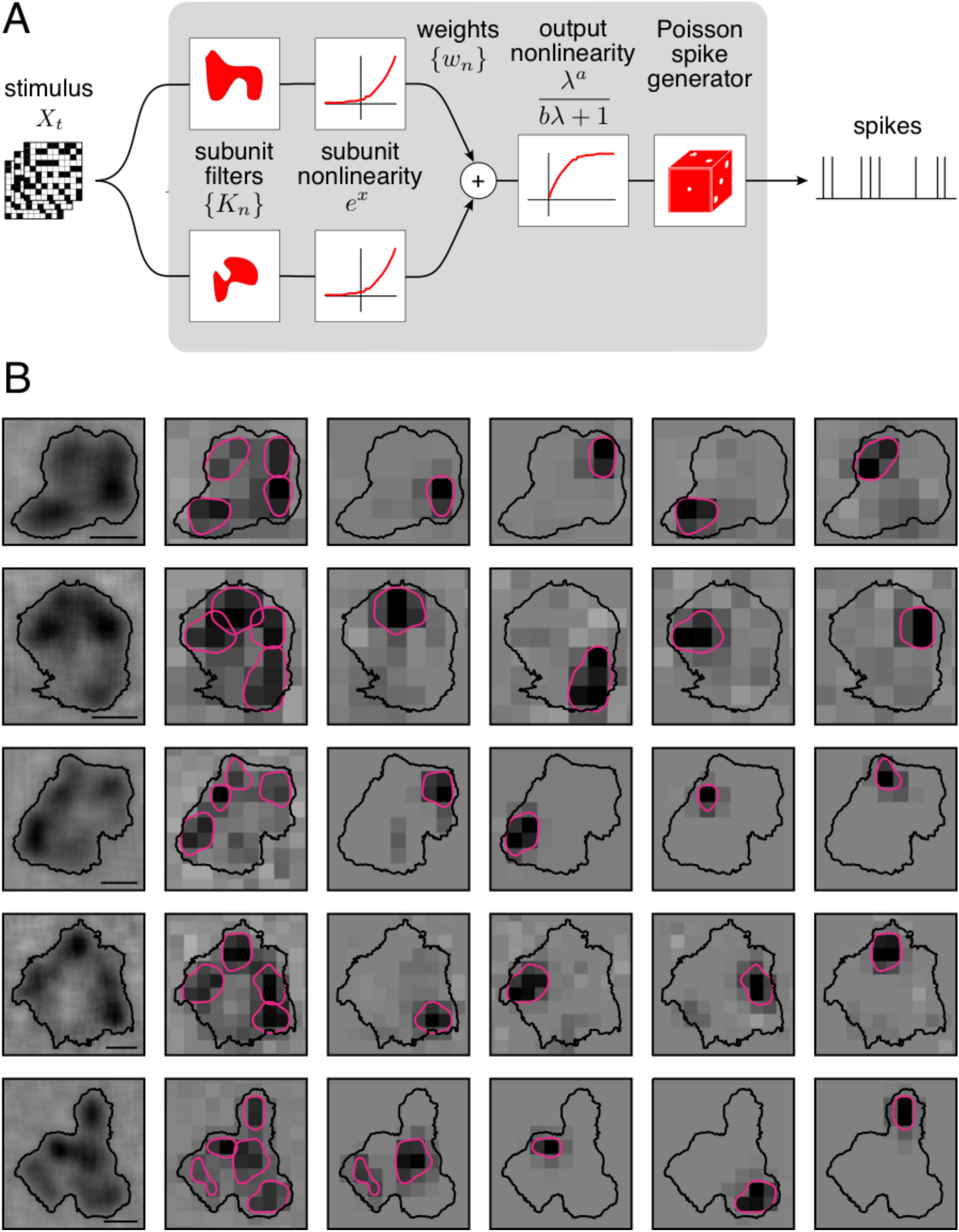
Subunit estimation through spike-triggered stimulus clustering. A: The model is constructed as a cascade of two LN stages. In the first stage, subunit activations were computed by linearly filtering the stimulus (X_t_) with kernels (K_n_) followed by an exponential nonlinearity. In the second stage, a sum of subunit activations, weighted by (w_n_), was passed through a saturating nonlinearity (g). Spikes were predicted with Poisson spike generation. B: Five example OFF SM cells were shown with the subunit model fits. Left: The RF of the SM cell, recorded on the jittering lattice, to observe hotspots. Contour threshold = 0.15. Second from left: The RF, recorded on a 84.8 × 84.8 μm coarse lattice for model fitting, and the contours of the subunit filters. Last four columns: the four subunit filters identified by the model as regions within which visual inputs sum linearly. The filters were Gaussian blurred, normalized, and contour fit with threshold = 0.4. Scale bars = 200 μm.

To test if the computational model identified the hotspots as subunits, the number of subunits was fixed at four. The regularization parameter was selected to maximize log-likelihood on a 10% validation dataset. The estimated subunits were then compared with the hotspots identified in the same cell. In cells that exhibited clear and distinct RF hotspots and for which model fits produced spatially localized subunit filters, a close alignment was frequently observed (Fig. 7B). Thus, RF hotspots represent strong sources of nonlinear computation in SM cells.

### Computational subunits align with regions with distinct spike waveforms

The previous results suggest a simple picture of SM cell visual computation: hotspots in the RF, formed in part by electrical compartments (Fig. 5), contribute to nonlinear computations over space (Figs. 6 and 7). A final test of consistency of this picture is to examine directly whether hotspots producing different spike waveforms align with the locations of computational subunits. A four-subunit model (Fig. 8) was fitted to the RF, and then compared to four compartments estimated from waveform clustering (Fig. 5B). For cells where four cleanly separated waveforms were observed, the computationally estimated subunits largely aligned with the regions identified by different spike waveforms, confirming their correspondence (Fig. 8A, B).

**Figure 8:**
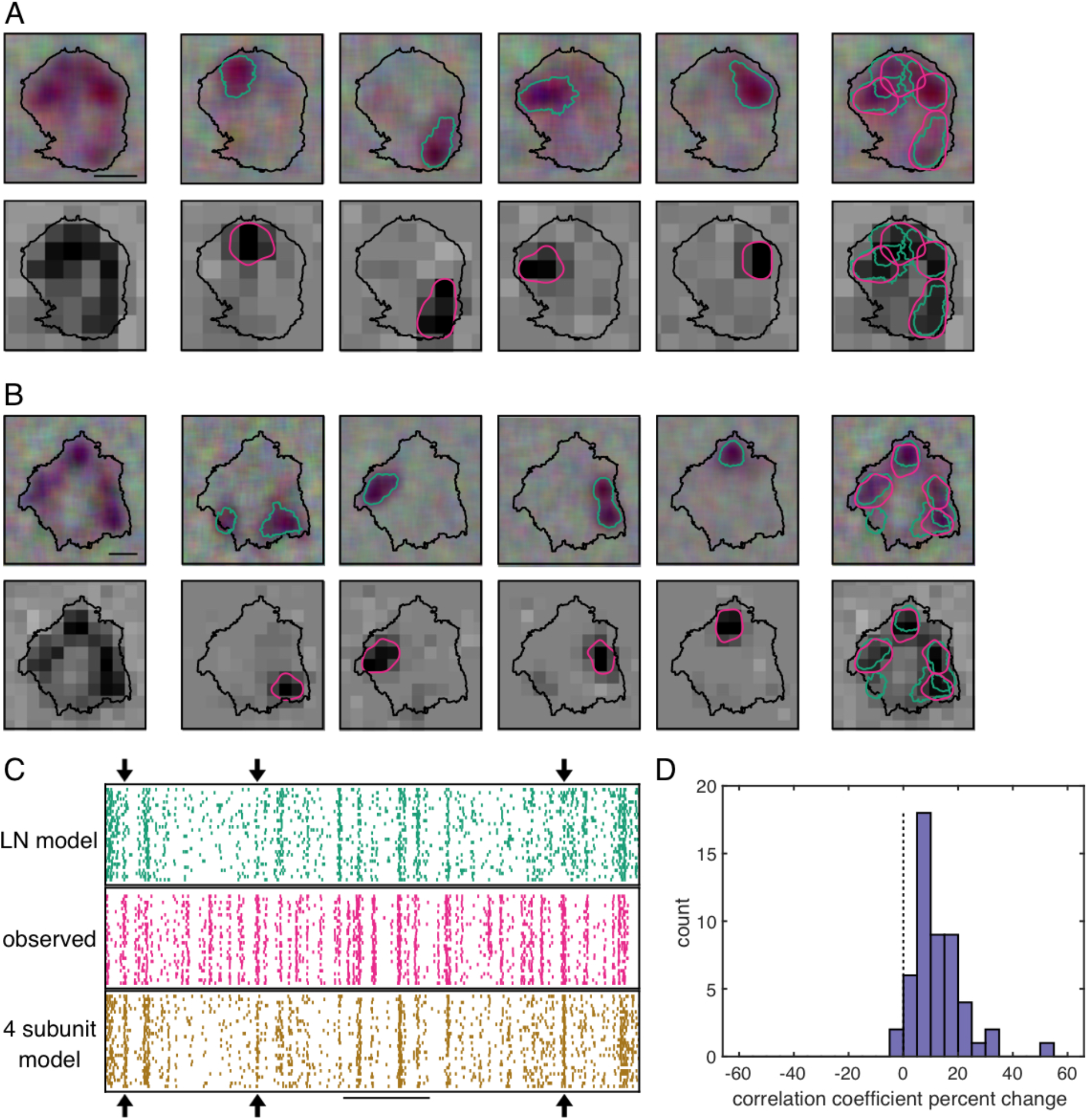
Computational subunits align with regions with distinct spike waveforms. A: Left column: The RF of an OFF SM recorded with white noise with a jittered lattice (top) or coarse white noise (bottom). Contour threshold = 0.15. Scale bar = 200 μm. Top middle four panels: The RF of four sub-clusters of distinct spike waveforms (contour threshold = 0.25) were identified as described previously (Fig. 5). Bottom middle four: the four subunit model filters, which were Gaussian blurred and contour fit at a threshold = 0.45. Right column: The correspondence of the teal spike waveform determined hotspots with the pink subunit model identified hotspots. B: Same as in A for another cell from a different recording. C: Middle: Observed responses of an SM cell to a 84.8 × 84.8 μm white noise stimulus for 30 repetitions (pink). Top: LN model prediction of the SM cell response. Bottom: Subunit model prediction of the SM cell response. Arrows indicate events captured by the subunit model that are not captured by the LN model. Scale bar = 1 second. D: Across 52 OFF SM cells from 10 recordings, the percent change in correlation coefficient of the subunit model compared to the LN model on a validation dataset consisting of 10% of the data.

To test if the subunit model resulted in more accurate predictions of SM cell responses than the commonly used LN model, predicted responses to repeated presentations of a white noise stimulus were computed for each model. At specific points during stimulus presentation, observed spikes were more accurately captured by the the subunit model than by the LN model (Fig 8C, arrows). Note, however, that many periods of firing were not accurately captured by either model. Therefore, while the subunit model resulted in more accurate predictions, more improvements to the model are necessary for a full understanding of SM cell function.

To compare the performance of the subunit model to a LN model across many cells and recordings, the correlation coefficient was computed on 10% held-out test data for the subunit and LN model. Across 52 OFF SM cells, there was a systematic increase in the correlation coefficient of the subunit model, compared to the LN model, which was reported as the percent change in correlation coefficient (12.6 ± 9.4%, mean ± SD across cells; Fig. 8D).

## Discussion

The physiological properties of the ON and OFF SM cells were investigated using large-scale multielectrode recordings, supplemented with patch clamp recordings to verify cell type identity. These two cell types exhibited complete mosaic coverage with precise interdigitation, as seen in the high-density RGC types. However, unlike the high-density RGC types, SM cells exhibited irregular RFs with distinct hotspots. The hotspots resulted in different spike waveforms when stimulated independently, suggesting unique biophysical mechanisms. A subunit computational model identified these hotspots as regions of nonlinear computation. The interpretation of these findings and the implications for understanding the diverse channels of visual processing emanating from the retina will be discussed.

### Non-traditional receptive field structure

The characteristic distinct hotspots observed in SM cell RFs have not been observed in other primate RGCs. Previously, canonical models such as a difference of Gaussian model of the RF (Enroth-Cugell and Robson, 1966; Rodieck and Stone, 1965), combined with the LN model of neural responses, have been used to describe RGCs (Chichilnisky, 2001). These models fail to adequately describe the structure (Fig. 3B) and responses (Fig. 8C, D) of the SM cells. First, the distinct hotspots in the RF depart strikingly from Gaussian models and motivate understanding the role of hotspots in computation. Second, the hotspots combine nonlinearly to drive firing, resulting in more predictive power from a computational model incorporating subunits than from an LN model. The nonlinearity could be produced entirely by bipolar cell rectification, as observed in other RGCs (Demb et al., 1999, 2001). However, alignment of the spike waveform differences and the strongest computational nonlinearities with hotspots (Fig. 8A) suggest an additional mechanism in SM cells, such as multiple electrical compartments.

### Identification of SM cells

Low-density primate RGC types have largely not been studied, due to the experimental challenges of recording spikes from rare cells. Large-scale MEA technology enabled recording complete populations of SM cells within a region of the retina (852 cells from 74 recordings of 48 macaque monkeys). However, variability across recordings, potentially arising from differences in the species, age, gender, health, and eccentricity, in addition to experimental considerations such as dissection technique and recording temperature, makes it difficult to identify SM cells reliably across preparations.

Normalizing time course properties by those of the easily identified, simultaneously recorded parasol cells revealed consistent physiological properties of identified SM cells in spite of this variability (Fig. 2). The narrow range of kinetic properties of putative SM cells corresponded closely with those of anatomically identified SM cells (Fig. 2B). This method relied on the assumption that among the low-density RGCs recorded using the present techniques, the recorded type with the fastest light response kinetics was usually the SM cells. This assumption is supported by the relatively large spike amplitude and size of the SM cell somas compared to other low-density cell types (Crook et al., 2008), making the SM cell spikes easier to detect and and segregate on the MEA. Potential failures of the approach are discussed further in Methods.

### Mechanism for spike waveform differences

The difference in extracellularly recorded spike waveforms elicited by stimulation of distinct hotspots supports the hypothesis that each hotspot has a distinct role in spike generation. One possibility is that spikes could be generated in the dendrites and propagated towards the soma. Another possibility is that synaptic currents carried by distinct bipolars to different regions of the dendrites could produce such an effect.

The number of distinct hotspots in each cell is approximately equal to the number of primary dendrites in SM cells (Crook et al., 2008; unpublished observations), suggesting that each hotspot could correspond to signals from a different primary dendrite. SM cell dendritic arbors differ from those of midget and parasol cells in at least two ways (Crook et al., 2008; Watanabe and Rodieck, 1989): 1) the SM cell primary dendrites leave the area near the soma uncovered and 2) distal dendrites are sparser (Dacey and Brace, 1992). These features could produce low sensitivity regions between dendritic branches, giving rise to hotspots in the RF. In addition, the nonlinear combination of presynaptic inputs could produce or amplify irregular RF structure produced by dendrites. Intracellular recordings combined with cell filling would be needed to test these hypotheses.

### Computational model of hotspot nonlinearity

Bipolar cell nonlinearities produce responses in RGCs to spatial patterns much finer than the RGC RF (Demb et al., 2001; Hochstein and Shapley, 1976; Victor and Shapley, 1979), and fine contrast reversing gratings have previously been used to understand the degree of nonlinearity in the inputs to different RGC types (Passaglia et al., 2002; Petrusca et al., 2007; Soodak et al., 1991). While observing RGC responses to fine stimuli confirms the presence of nonlinear computation, understanding the specific nonlinear components of each cell, their arrangement over space, and their particular role in visual signaling requires additional tests.

Nonlinear signaling in SM cells was probed with two methods. In a closed-loop experiment, RF hotspots were independently visually stimulated. This revealed strong firing when one spot strongly drives the cell and the other provides a suppressive input, consistent with a strong nonlinearity associated with hotspots (Fig. 6). While this result could be explained simply by rectifying nonlinearities in bipolar cells composing the hotspots, the model confirms that the hotspots are important for the nonlinear visual computations performed in SM cells.

In addition, the computational model identified spatially localized regions that aligned with the hotspots (Fig. 8). This model explained SM cell responses more accurately than the commonly used LN models, which do not account for spatial nonlinearities. However, much is still unknown about the connectivity and nonlinearities in the SM cell circuit. For example, it is not clear how the hotspots could relate to bipolar cells providing input to SM cells: bipolar cell locations are not easily measured, but the known sizes of the bipolar cells presynaptic to the SM cells suggests that each hotspot would probably be composed of several bipolar cells, possibly of multiple types (Boycott and Wassle, 1991; Crook et al., 2008; Dacey et al., 2000; Jacoby et al., 2000; Tsukamoto and Omi, 2015, 2016). In the future, a more complete model based on additional information about the underlying circuitry could potentially predict the responses more accurately.

### The role of SM cells in vision

The primate retina has ~20 parallel visual channels, each of which extracts different information about the external world from the visual image. The function of the five most common RGC types in primate vision has been investigated extensively, but the role of most of the remaining ~15 cell types is largely unknown. In other mammals, like rodents, rabbits, and cats, specialized functions such as direction selectivity (Barlow et al., 1964), object motion selectivity (Olveczky et al., 2003), local edge detection (Levick, 1967), looming detection (Münch et al., 2009), anticipation of motion (Berry et al., 1999), and suppression by motion (Tien et al., 2015) have been observed. The irregular hotspot structure and nonlinear computation in SM cells suggest a mechanism for visual signalling not previously identified in the primate retina, but the precise role of these cells in visual signaling remains an open question.

## Acknowledgements

This work was supported by NIH NEI F31EY027166 (CR), NSF GRFP DGE-114747 (CR, NB), NIH NEI R01-EY027323 (MM), NIH NEI P30-EY001730 (UW Vision Core; MM, Research to Prevent Blindness Unrestricted Grant (UW Department of Ophthalmology; MM), NSF IGERT Grant 0801700 (NB), Wu Tsai Neurosciences Institute Interdisciplinary Scholar Award (GG), Pew Charitable Trust Scholarship in the Biomedical Sciences (AS), NIH NEI R01-EY021271 (EJC), NIH NEI P30-EY019005 (EJC), NIH NEI R01-EY029247 (MM and EJC). We thank Fred Rieke for helpful suggestions on the manuscript, Jill Desnoyer and Ryan Samarakoon for technical assistance, and H. Fox, M. Taffe, T. Albright, R. Krausliz, R. Siegel, K. Bankiewicz, C. Darian-Smith, J. Carmena, J. Wallis, E. Callaway, T. Moore, S. Morairty, the UC Davis Primate Center, and Washington National Primate Research Center (NIH P51 OD-010425) for providing access to retinas.

## Author contributions

CR, NS, NB, AK, GG collected the MEA data, MM collected and analyzed the patch clamp data, CR, EJC conceived and designed the experiments, CR analyzed the data, NS developed the subunit model, GG developed analysis software, AS and AL developed and supported the MEA hardware and software, CR and EJC wrote the paper, all authors edited the paper, EJC supervised the project.

## Methods

### Electrophysiology and anatomical verification

Large-scale multi-electrode recordings were performed from retinas of macaque monkeys, as described previously (Chichilnisky and Baylor, 1999; Field et al., 2007). Briefly, eyes were removed from terminally anesthetized macaque monkeys (*Macaca mulatta, Macaca fascicularis*) used by other laboratories in the course of their experiments, in accordance with the Institutional Animal Care and Use Committee guidelines. Prior to enucleation, the animals underwent various procedures including SIV infections, rabies injections, and chronic cortical recording implants. Some animals suffered from health problems including diabetes, arthritis, dementia, seizures, or hepatic amyloidosis but no visual impairments were known. Following enucleation, the anterior portion of the eye and vitreous were removed. The eye was stored in a dark container in oxygenated Ames’ solution (Sigma, St. Louis, MO) at 33°C, pH 7.4. Under infrared illumination, a small piece of retina approximately 3×3 mm, from a retinal region with eccentricity between 4.5-17 mm (4.0-17 mm temporal equivalent eccentricity (Chichilnisky and Kalmar, 2002) or 20-82.6 degrees (Dacey and Petersen, 1992; Perry and Cowey, 1985) was dissected and placed ganglion cell side down on a multi-electrode array for recording. In some preparations, the RPE remained attached during the recording; in others the RPE was removed. For the RPE attached recordings, the choroid was removed up to Bruch’s membrane to improve oxygenation and maintain even retinal thickness. RPE-attached recordings were performed at 30-32°C and isolated retina recordings at 32-37°C. For the duration of the recording, the retina was perfused with oxygenated Ames’ solution.

Several types of custom planar large-scale multielectrode arrays were used. The array either had a rectangular outline with 16 × 32 electrodes at 60 μm pitch or a hexagonal outline with 519 electrodes at 120 μm or 30 μm pitch (Field et al., 2010; Litke et al., 2004). Recorded voltages were bandpass filtered, amplified, and digitized at 20 kHz using custom electronics (Litke et al., 2004). Recordings were analyzed to isolate spikes produced by different cells, a process called spike sorting, summarized briefly here (Field et al., 2007). To detect spikes on each electrode, a threshold of four times the standard deviation of the voltage over time was used. Voltage waveforms surrounding the time of the spike (−0.5 to 0.75 ms) on that electrode and the surrounding six electrodes were extracted and concatenated into a vector. Principal component analysis, typically followed by noise whitening, and unsupervised clustering using a mixtures of Gaussian model on the first five principal components of the vector waveforms was performed (Duda et al., 2001). Clusters consisting of more than 100 spikes were identified as neurons if a 1 ms refractory period was observed among the spikes in that cluster. Duplicate copies of neurons detected by multiple electrodes were identified by temporal cross-correlation and were removed. SM cell spikes were more difficult to accurately cluster than those of other RGC types because of the lower firing rate and the smaller spike amplitude. In addition to the automated clustering described above, the clusters on every electrode where an SM cell was identified were manually verified. If necessary, clusters were merged or split if incorrectly identified by automated clustering.

For the patch clamp experiments, recordings were performed from an RPE-attached retinal preparation at eccentricities of 4-8 mm (4-8 mm temporally equivalent eccentricity or 18-36 degrees) in three macaque retinas. Recordings were performed using borosilicate glass pipettes as described previously (Manookin et al., 2015). Cells were filled with biocytin (0.5%; EZ-Link, ThermoFisher) for later anatomical imaging. Following recording, tissue was immersion fixed in a solution containing 4% PFA for 30-45 min and washed with 1X PBS. Tissue was counterstained with Streptavidin-conjugated Alexa 488 to visualize filled cells. Images of filled cells were acquired using a SP8 confocal microscope (Leica) and used for anatomical verification.

### Visual stimulation

The image from a gamma-corrected CRT monitor (Sony Trinitron Multiscan E100; Sony, Tokyo, Japan) refreshing at 120 Hz was optically reduced and projected onto the retina. In RPE attached recordings, the visual stimulus was delivered through the mostly-transparent electrode array. In isolated retina recordings, the visual stimulus was delivered from the photoreceptor side. For all experiments except those in Fig. 2, low photopic light levels were used (rates of 800-2200, 800-2200, and 400-900 photoisomerizations per second for the L, M and S cones respectively; see Field et al., 2009, 2010). For Fig. 2, the light levels spanned the photopic and scotopic ranges.

A white noise stimulus consisting of a lattice of pixels flickering independently at 15-120Hz was used to characterize the spatial, temporal, and chromatic response properties of the recorded RGCs (Chichilnisky, 2001). Different experiments utilized different pixel sizes, ranging from 21.2 to 106 μm on a side. On each refresh, the intensity of each display primary at each pixel location was assigned a value randomly selected from a binary distribution. The pixel contrast (standard deviation of the difference from the mean pixel intensity) was 96% for each display primary. In some experiments, 84.8 μm × 84.8 μm pixels were displayed on a underlying lattice jittering on a 5.3 μm grid. The duration of each recording with white noise stimulation was 15-300 min.

### Light response characterization

The spike triggered average (STA) stimulus for each neuron was computed from the response to the white noise stimulus (Chichilnisky, 2001), to reveal spatial, temporal, and chromatic properties of the light response.

Previously, 2D Gaussian fits have been used to characterize the RF size of RGCs (Enroth-Cugell and Robson, 1966; Enroth-Cugell et al., 1983; Rodieck and Stone, 1965). However, the RFs of the SM cells have distinct hotspots (Fig. 3B) and thus are not accurately summarized with such a fit (Brown et al., 2000; Gauthier et al., 2009; Rowe and Cox, 1993; Rowe and Palmer, 1995; Schwartz et al., 2012; Soo et al., 2011; Soodak et al., 1991). For each SM cell, a contour was used to summarize the RF. The threshold of the contour, after normalizing the RF to a maximum amplitude of 1, varied due to differences in SNR among the SM cell recordings and varying degrees of separation of the hotspots. The threshold used in each figure panel is reported in the caption. For Fig. 5G-H, the convex hull of the contour was shown due to the separation of the hotspots.

In Fig. 1, to visualize the mosaic coverage typically reported using difference of Gaussians fitting (Devries and Baylor, 1997), the contours were shrunk by 30% around the center. The shrinkage was applied because the white noise stimulus was displayed on a underlying jittering lattice that resulted in spatial correlations that artificially extended the size of the RF contour. A higher contour threshold was not used because it failed to capture the complete RF of the SM cells with strong hotspot structure. The contours in Fig. 1 should not be interpreted as the actual RF sizes, but rather the relative sizes of the various RGC types.

The response time course (Fig. 1, 2) was obtained by averaging the values of the significant pixels in the STA across time. Significant pixels were defined as those with an absolute maximum value more than 4.5 times the robust standard deviation (Freeman et al., 2015) of all the pixels in the STA. The RF was visualized as the frame of the STA corresponding to the peak of the response time course. The autocorrelation functions (Fig. 1) were normalized to unit area.

### Cell type classification

Cell type classification within a recording was performed by identifying distinct clusters in the response properties of all cells recorded with white noise stimulation (Fig. 1; Field et al., 2007). In Fig. 1, the area of the RF contour for each cell and the first principal component of the STA time courses were examined for classification. In other recordings, the second principal component of the time course and the principal components of the autocorrelation functions were also used (not shown). Neurons with evidence of contamination of spikes from another cell, low spike counts, or low SNR were removed from consideration. The similarities of autocorrelation functions within a cell type and the differences across cell types also confirmed cell type classification (Fig. 1; Devries and Baylor, 1997). In each recording, the correspondence between functionally recorded RGC types and anatomically identified types for ON parasol, OFF parasol, ON midget, OFF midget, and small bistratified cells were inferred from cell densities and light response properties (Chichilnisky and Kalmar, 2002; Field et al., 2007; Dacey 2004). All other RGCs were identified as low-density cells.

To classify the distinct types of low-density cells, a collection of 53 recordings, each containing multiple ON and/or multiple OFF low-density cells were examined. For each recording, cell type classification was performed (Fig. 1) based on RF size, time course, autocorrelation function, and mosaic overlap. The low-density cell type with the fastest kinetics in the recording, measured by the time of zero crossing (Fig. 2A) was labeled α, and the other low-density cell types were combined and labeled β.

To determine if α cells had reproducible kinetic properties across recordings, the time of zero crossing and biphasic index were computed with the following procedure: 1) the time course was averaged from the significant pixels and smoothed with a moving average of span two. 2) The time of zero crossing was linearly interpolated from the measurements around zero. 3) The biphasic index was measured as the maximum absolute amplitude of the signal in the peak lobe divided by the maximum absolute amplitude of the signal in the trough lobe. 4) The time of zero crossing and biphasic index of ON (OFF) low-density cells were normalized by the same measurement from the simultaneously recorded population average of the ON (OFF) parasol cells.

The α and β cells were distinct based on these two normalized metrics. To determine the boundary between the α and β cell clusters, a support vector machine was fit with a radial kernel (Boser et al., 1992). The classification boundary was selected to contain all points that the support vector machine classified as α with greater than 95% probability. To confirm the anatomical identity of the α cells, the same parameters were computed on morphologically identified SM cells with single cell patch clamp (Fig. 2B).

The success of the approach is supported by considering what would happen under two potential failure modes: 1) no SM cells were recorded and 2) the classification boundary included multiple cell types. They will be addressed in turn:

1) If SM cells were not recorded in a particular preparation, but multiple other types of low-density cells were recorded, the cells labeled α would not be SM cells, and would have time course properties diverging from the majority of the α cluster. These cells would presumably appear as outliers from the tightly clustered α points. As long as the majority of recordings did contain SM cells, the conservatively defined classification boundary would likely not be affected greatly by these outliers. Indeed, some recordings contained no cells within the classification boundary.
2) If there were two or more low-density cell types with similar kinetic properties, then across 53 recordings it is likely that two or more of those cell types would be recorded in the same retina, revealing failures of mosaic organization. In this case, α and β would have similar kinetic properties. Instead, however, in no cases did both α and β appear within the SM cell classification boundary. This observation suggests that the boundary is restrictive enough to exclude other RGC types.

### Photoreceptor control analysis

For each OFF SM cell, the contour fit at a threshold of 0.15 on the RF normalized to a peak of 1 was computed. The OFF parasol cells whose RF centers (defined by a 2D Gaussian fit) fell within the convex hull of the contour fit of the SM cell were selected for analysis. The SNR of each parasol cell was computed by dividing the peak value of the Gaussian fit by the standard deviation of a 10×10 background pixel region (each pixel is 42.8 × 42.8 μm). If the average of the SM cell RF within the parasol cell RF, defined by 1 standard deviation of Gaussian fit, was less than a threshold, then the parasol cell was defined as “strong”; otherwise, the parasol cell was classified as “weak”. The mean ± SEM of the SNR for the strong and the weak parasol cells in each recording were shown (Fig. 3D). OFF SM cells were exclusively used for this control because the OFF SM cells exhibited more separated and distinct hotspots than the ON SM cells (see Fig. 3B).

### Interdigitation analysis

For analysis of interdigitation (Fig. 4A-D), mosaics of five or more neighboring ON or OFF SM cells recorded with white noise (pixel size 21.2 - 53.0 μm) were examined. The RF was blurred with a Gaussian (SD = 21.2 μm) before fitting a contour to the spatial RF. For each cell type in each recording, the same contour threshold was applied to all cells. The optimal contour threshold was defined as the value that maximized the UI: the number of pixels covered by the contour of exactly one cell, divided by all of the pixels within the bounded region. The boundary of the region was determined by performing the procedure outlined previously (Gauthier et al., 2009). Briefly, a Delaunay Triangulation of the centers of the contour fits of all cells was computed. The area within a triangle was included in the overall boundary if the cells at its vertices were closer than 1.9 times the median nearest neighbor spacing of all the cells in the mosaic, indicating no apparent gap in the mosaic. The UI was computed within the boundary.

The UI was computed for the observed data as well as 1000 mosaics obtained by random rotations of the measured RFs. The RF of each cell was rotated around the center of the contour fit. The contour threshold was then recomputed, maximizing the UI. The distribution of UI values produced by the 1000 perturbations (Fig. 4C) was treated as the null distribution for statistical testing.

### Shared input

To determine which cell pairs had overlapping hotspots, strong pixels (with an intensity greater than 50% of the maximum value) were identified for each cell. If the two cells shared strong pixels, then the cell pair was identified as sharing a hotspot. A square was identified, centered at the pixel with the maximum intensity product in the two RFs. The correlation coefficient between the pixel intensities in the two cells within the square was computed for the observed data, as well as for 7 manipulations in which the data from one cell was rotated by 0, 90, 180, or 270 degrees, and/or flipped horizontally around the center point. The coefficients were ranked in descending order and the rank of the observed data compared to the various manipulations was reported, where rank 1 indicated highest correlation. If the hotspots in the two cells had identical sensitivity profiles, then the observed data would have a rank of 1.

### Spike waveform splitting

In the spike sorting procedure (see above), sub-clusters were sometimes observed within the cluster corresponding to a single SM cell. An automated procedure was developed to identify such sub-clusters. For each electrode, the standard spike sorting approach as described previously was applied. Then 50 iterations of spectral clustering (Ng et al., 2001; Zelnik-Manor and Perona, 2004) with random initial conditions were performed on the first five principal components of the spike waveforms from the SM cell with the number of clusters equal to the number of visible hotspots in the RF. The iteration with the smallest average difference from the rest of the iterations in spike-to-cluster assignments was selected. In general less than 5% of spike waveform cluster assignments varied among iterations of spectral clustering. The RFs of each cluster were then computed, fitted with contours, and compared to the RF computed from all the SM cell spikes. The threshold for the contour fit of the SM cells and the hotspots in this analysis was 0.2.

The EI for each cluster on the electrode was computed as well as the EI for the subclusters. For every spike of a RGC, the voltage waveform 2.5 ms before to 2.5 ms after was averaged for every electrode on the array. By averaging across all of the spikes of a neuron and taking the maximum projection across time, the EI revealed the unique somatic, dendritic, and axonal signatures of each neuron. The outline of the maximum projection of the EI (Fig. 5G, H), was the compact boundary around the electrodes in the combined cluster where the voltage fell below −30 μV.

To observe the spatial separation in the origin of electrical signal, the automated waveform clustering analysis was performed on SM cells recorded on an array with 30 μm spacing. The time preceding the spike at which the waveform first fell below a threshold of 5.5 μV was recorded for each electrode and each cluster. The threshold was chosen to be robust to noise (Fig. 5J: green asterisks). This measured time on each electrode for the second cluster (Fig. 5H: dotted line) was subtracted from the time for the first cluster (Fig. 5H: solid line). Red electrodes in Fig. 5I were those for which the first cluster was faster than the second cluster, and vice versa for blue electrodes. Electrodes for which the difference was less than 50 μV were not shown.

### Closed loop white noise stimulus

For closed loop visual stimulation (Fig. 6), the STA was computed following spike sorting while the recording was taking place. Two circular stimulation spots were placed to overlap with the hotspots observed in the RF. The spot size was adjusted so that both encompassed approximately the same integrated weight of the STA. Each spot was independently stimulated with binary white noise for 15-30 minutes. The temporally integrated contrast on each spot was determined by convolving the stimulus with the time course of the cell. The firing rate was determined by counting spikes in 8.33 ms bins corresponding to stimulus display frames. The effective contrast values on both spots, as well as the firing rate, were collected into 10×10×10 bins of variable size such that each bin contained the same number of samples. The linear model is described by:

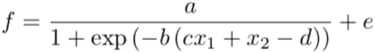

The nonlinear model is described by:

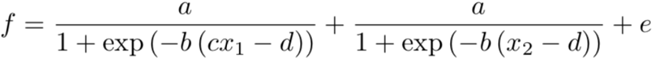

where x_1_ and x_2_ are the binned input contrast values, and a, b, c, d, and e are the five fit parameters. The models were fitted for each cell minimizing the mean squared error between f and the recorded binned firing rate.

### Spiking response model fitting

The subunit model construction and fitting approach were described in Shah et al., 2018. Briefly, the model was fit using an alternating cascade of two linear-nonlinear stages with a Poisson firing rate. In the model, the visual stimulus was convolved with the average time course of the neuron to produce the instantaneous stimulus (X_t_). The inner product of the stimulus with each of the model linear subunit filters (K_n_) was computed (where the total number of desired subunits was fixed), and followed by an exponential nonlinearity. Then, the subunit outputs were weighted by non-negative weights (w_n_), summed, and passed through a final output nonlinearity of the form g(x) = x^a / (bx+1), resulting in the firing rate: g(Σw_n_ exp(K_n_•X_t_)). Spikes were predicted from this rate with a Poisson spike generator. A locally normalized L1 regularizer was used which penalizes large weights if all of the neighboring weights are small. The model was fitted on 80% of the data.

For simplicity, the number of subunit filters was fixed at four in order to approximately correspond to the number of visible hotspots. A range of regularization values was searched for each cell, and the value that maximized log likelihood on a 10% test dataset were used.

The LN model, used for comparison to the performance of the subunit model (Fig. 8), was fit using the same method previously described for the subunit model, except the number of subunits was fixed at one. The reported percentage improvement in correlation coefficient was measured as the difference between the correlation coefficient of the four subunit model and the LN model divided the correlation coefficient of the LN model. The correlation coefficient was measured on three minutes of white noise data not used in either the fitting or model selection process.

---

**Supplemental Movie 1: Dendritic origin of waveform differences**

The electrical image over time of the two distinct SM cell waveform clusters shown in Fig. 5H. The electrical signal corresponding to different hotspots in the RF originates from different spatial locations, as shown at approximately −250 μs (and summarized in Fig. 5I). Each black dot is an electrode on the 519 hexagonal multi-electrode array. Yellow indicates negative voltage deflection while purple indicates positive voltage deflection. The size indicates the magnitude of the voltage deflection. At 0 μs, the spike occurs. The red dots mark the electrode with maximal overall deflection (approximately the soma location).

## References

Barlow, H.B., and Levick, W.R. (1965). The mechanism of directionally selective units in rabbit’s retina. J. Physiol. 178, 477–504.

Barlow, H.B., Hill, R.M., and Levick, W.R. (1964). RETINAL GANGLION CELLS RESPONDING SELECTIVELY TO DIRECTION AND SPEED OF IMAGE MOTION IN THE RABBIT. J. Physiol. 173, 377–407.

Berry, M.J., 2nd, Brivanlou, I.H., Jordan, T.A., and Meister, M. (1999). Anticipation of moving stimuli by the retina. Nature 398, 334–338.

Boser, B.E., Guyon, I.M., and Vapnik, V.N. (1992). A training algorithm for optimal margin classifiers. In Proceedings of the Fifth Annual Workshop on Computational Learning Theory - COLT’ 92,.

Boycott, B.B., and Wassle, H. (1991). Morphological Classification of Bipolar Cells of the Primate Retina. Eur. J. Neurosci. 3, 1069–1088.

Brown, S.P., He, S., and Masland, R.H. (2000). Receptive field microstructure and dendritic geometry of retinal ganglion cells. Neuron 27, 371–383.

Chichilnisky, E.J. (2001). A simple white noise analysis of neuronal light responses. Network 12, 199–213.

Chichilnisky, E.J., and Baylor, D.A. (1999). Receptive-field microstructure of blue-yellow ganglion cells in primate retina. Nat. Neurosci. 2, 889–893.

Chichilnisky, E.J., and Kalmar, R.S. (2002). Functional asymmetries in ON and OFF ganglion cells of primate retina. J. Neurosci. 22, 2737–2747.

Crook, J.D., Peterson, B.B., Packer, O.S., Robinson, F.R., Gamlin, P.D., Troy, J.B., and Dacey, D.M. (2008). The smooth monostratified ganglion cell: evidence for spatial diversity in the Y-cell pathway to the lateral geniculate nucleus and superior colliculus in the macaque monkey. J. Neurosci. 28, 12654–12671.

Dacey, D.M. (1993a). The mosaic of midget ganglion cells in the human retina. J. Neurosci. 13, 5334–5355.

Dacey, D.M. (1993b). Morphology of a small-field bistratified ganglion cell type in the macaque and human retina. Vis. Neurosci. 10, 1081–1098.

Dacey, D.M. (2004). Origins of perception: retinal ganglion cell diversity and the creation of parallel visual pathways. In The cognitive neurosciences III, Gazzaniga MS, ed (Cambridge, MA: MIT), pp 281–301.

Dacey, D.M., and Brace, S. (1992). A coupled network for parasol but not midget ganglion cells in the primate retina. Vis. Neurosci. 9, 279–290.

Dacey, D.M., and Petersen, M.R. (1992). Dendritic field size and morphology of midget and parasol ganglion cells of the human retina. Proc. Natl. Acad. Sci. U. S. A. 89, 9666–9670.

Dacey, D., Packer, O.S., Diller, L., Brainard, D., Peterson, B., and Lee, B. (2000). Center surround receptive field structure of cone bipolar cells in primate retina. Vision Res. 40, 1801–1811.

Dacey, D.M., Peterson, B.B., Robinson, F.R., and Gamlin, P.D. (2003). Fireworks in the Primate Retina. Neuron 37, 15–27.

Dacey, D.M., Liao, H.-W., Peterson, B.B., Robinson, F.R., Smith, V.C., Pokorny, J., Yau, K.-W., and Gamlin, P.D. (2005). Melanopsin-expressing ganglion cells in primate retina signal colour and irradiance and project to the LGN. Nature 433, 749–754.

Demb, J.B., Haarsma, L., Freed, M.A., and Sterling, P. (1999). Functional circuitry of the retinal ganglion cell’s nonlinear receptive field. J. Neurosci. 19, 9756–9767.

Demb, J.B., Zaghloul, K., Haarsma, L., and Sterling, P. (2001). Bipolar cells contribute to nonlinear spatial summation in the brisk-transient (Y) ganglion cell in mammalian retina. J. Neurosci. 21, 7447–7454.

Devries, S.H., and Baylor, D.A. (1997). Mosaic arrangement of ganglion cell receptive fields in rabbit retina. J. Neurophysiol. 78, 2048–2060.

Duda, R.O., Hart, P.E., and Stork, D.G. (2001). Pattern classification. (New York: Wiley).

Enroth-Cugell, C., and Robson, J.G. (1966). The contrast sensitivity of retinal ganglion cells of the cat. J. Physiol. 187, 517–552.

Enroth-Cugell, C., Robson, J.G., Schweitzer-Tong, D.E., and Watson, A.B. (1983). Spatio-temporal interactions in cat retinal ganglion cells showing linear spatial summation. J. Physiol. 341, 279–307.

Field, G.D., Sher, A., Gauthier, J.L., Greschner, M., Shlens, J., Litke, A.M., and Chichilnisky, E.J. (2007). Spatial properties and functional organization of small bistratified ganglion cells in primate retina. J. Neurosci. 27, 13261–13272.

Field, G.D., Greschner, M., Gauthier, J.L., Rangel, C., Shlens, J., Sher, A., Marshak, D.W., Litke, A.M., and Chichilnisky, E.J. (2009). High-sensitivity rod photoreceptor input to the blue-yellow color opponent pathway in macaque retina. Nat. Neurosci. 12, 1159–1164.

Field, G.D., Gauthier, J.L., Sher, A., Greschner, M., Machado, T.A., Jepson, L.H., Shlens, J., Gunning, D.E., Mathieson, K., Dabrowski, W., et al. (2010). Functional connectivity in the retina at the resolution of photoreceptors. Nature 467, 673–677.

Frechette, E.S., Sher, A., Grivich, M.I., Petrusca, D., Litke, A.M., and Chichilnisky, E.J. (2005). Fidelity of the ensemble code for visual motion in primate retina. J. Neurophysiol. 94, 119–135.

Freeman, J., Field, G.D., Li, P.H., Greschner, M., Gunning, D.E., Mathieson, K., Sher, A., Litke, A.M., Paninski, L., Simoncelli, E.P., et al. (2015). Mapping nonlinear receptive field structure in primate retina at single cone resolution. Elife 4.

Gauthier, J.L., Field, G.D., Sher, A., Greschner, M., Shlens, J., Litke, A.M., and Chichilnisky, E.J. (2009). Receptive fields in primate retina are coordinated to sample visual space more uniformly. PLoS Biol. 7, e1000063.

Gollisch, T., and Meister, M. (2010). Eye smarter than scientists believed: neural computations in circuits of the retina. Neuron 65, 150–164.

Greschner, M., Field, G.D., Li, P.H., Schiff, M.L., Gauthier, J.L., Ahn, D., Sher, A., Litke, A.M., and Chichilnisky, E.J. (2014). A polyaxonal amacrine cell population in the primate retina. J. Neurosci. 34, 3597–3606.

Hochstein, S., and Shapley, R.M. (1976). Linear and nonlinear spatial subunits in Y cat retinal ganglion cells. J. Physiol. 262, 265–284.

Jacoby, R.A., Wiechmann, A.F., Amara, S.G., Leighton, B.H., and Marshak, D.W. (2000). Diffuse bipolar cells provide input to OFF parasol ganglion cells in the macaque retina. J. Comp. Neurol. 416, 6–18.

Kuffler, S.W. (1953). Discharge patterns and functional organization of mammalian retina. J. Neurophysiol. 16, 37–68.

Levick, W.R. (1967). Receptive fields and trigger features of ganglion cells in the visual streak of the rabbit’s retina. J. Physiol. 188, 285–307.

Li, P.H., Gauthier, J.L., Schiff, M., Sher, A., Ahn, D., Field, G.D., Greschner, M., Callaway, E.M., Litke, A.M., and Chichilnisky, E.J. (2015). Anatomical identification of extracellularly recorded cells in large-scale multielectrode recordings. J. Neurosci. 35, 4663–4675.

Liao, H.-W., Ren, X., Peterson, B.B., Marshak, D.W., Yau, K.-W., Gamlin, P.D., and Dacey, D.M. (2016). Melanopsin-expressing ganglion cells on macaque and human retinas form two morphologically distinct populations. J. Comp. Neurol. 524, 2845–2872.

Litke, A.M., Bezayiff, N., Chichilnisky, E.J., Cunningham, W., Dabrowski, W., Grillo, A.A., Grivich, M., Grybos, P., Hottowy, P., Kachiguine, S., et al. (2004). What does the eye tell the brain?: Development of a system for the large-scale recording of retinal output activity. IEEE Trans. Nucl. Sci. 51, 1434–1440.

Liu, J.K., Schreyer, H.M., Onken, A., Rozenblit, F., Khani, M.H., Krishnamoorthy, V., Panzeri, S., and Gollisch, T. (2017). Inference of neuronal functional circuitry with spike-triggered nonnegative matrix factorization. Nat. Commun. 8, 149.

Manookin, M.B., Puller, C., Rieke, F., Neitz, J., and Neitz, M. (2015). Distinctive receptive field and physiological properties of a wide-field amacrine cell in the macaque monkey retina. J. Neurophysiol. 114, 1606–1616.

Manookin, M.B., Patterson, S.S., and Linehan, C.M. (2018). Neural Mechanisms Mediating Motion Sensitivity in Parasol Ganglion Cells of the Primate Retina. Neuron 97, 1327–1340.e4.

Masland, R. (2001). Neuronal diversity in the retina. Curr. Opin. Neurobiol. 11, 431–436.

Münch, T.A., da Silveira, R.A., Siegert, S., Viney, T.J., Awatramani, G.B., and Roska, B. (2009). Approach sensitivity in the retina processed by a multifunctional neural circuit. Nat. Neurosci. 12, 1308–1316.

Ng, A., Jordan, M., and Weiss, Y. (2001). On Spectral Clustering: Analysis and an algorithm. 14th Conference of Neural Information Processing Systems.

Olveczky, B.P., Baccus, S.A., and Meister, M. (2003). Segregation of object and background motion in the retina. Nature 423, 401–408.

Passaglia, C.L., Troy, J.B., Rüttiger, L., and Lee, B.B. (2002). Orientation sensitivity of ganglion cells in primate retina. Vision Res. 42, 683–694.

Peichl, L. (1991). Alpha ganglion cells in mammalian retinae: common properties, species differences, and some comments on other ganglion cells. Vis. Neurosci. 7, 155–169.

Perry, V.H., and Cowey, A. (1985). The ganglion cell and cone distributions in the monkey’s retina: implications for central magnification factors. Vision Res. 25, 1795–1810.

Petrusca, D., Grivich, M.I., Sher, A., Field, G.D., Gauthier, J.L., Greschner, M., Shlens, J., Chichilnisky, E.J., and Litke, A.M. (2007). Identification and characterization of a Y-like primate retinal ganglion cell type. J. Neurosci. 27, 11019–11027.

Puller, C., Manookin, M.B., Neitz, J., Rieke, F., and Neitz, M. (2015). Broad thorny ganglion cells: a candidate for visual pursuit error signaling in the primate retina. J. Neurosci. 35, 5397–5408.

Ravi, S., Ahn, D., Greschner, M., Chichilnisky, E.J., and Field, G.D. (2018). Pathway-specific asymmetries between ON and OFF visual signals. J. Neurosci.

Rodieck, R.W. (1998). The First Steps in Seeing (Sinauer Associates Incorporated).

Rodieck, R.W., and Stone, J. (1965). ANALYSIS OF RECEPTIVE FIELDS OF CAT RETINAL GANGLION CELLS. J. Neurophysiol. 28, 833–849.

Roska, B., and Meister, M. (2014). The retina dissects the visual scene into distinct features. In The new visual neurosciences, J.H. Werner & L.M. Chalupa, eds (Cambridge, MA: MIT Press), pp. 163–183.

Rowe, M.H., and Cox, J.F. (1993). Spatial receptive-field structure of cat retinal W cells. Vis. Neurosci. 10, 765–779.

Rowe, M.H., and Palmer, L.A. (1995). Spatio-temporal receptive-field structure of phasic W cells in the cat retina. Vis. Neurosci. 12, 117–139.

Schwartz, G.W., Okawa, H., Dunn, F.A., Morgan, J.L., Kerschensteiner, D., Wong, R.O., and Rieke, F. (2012). The spatial structure of a nonlinear receptive field. Nat. Neurosci. 15, 1572–1580.

Shapley, R.M., and Victor, J.D. (1979). Nonlinear spatial summation and the contrast gain control of cat retinal ganglion cells. J. Physiol. 290, 141–161.

Soo, F.S., Schwartz, G.W., Sadeghi, K., and Berry, M.J., 2nd (2011). Fine spatial information represented in a population of retinal ganglion cells. J. Neurosci. 31, 2145–2155.

Soodak, R.E., Shapley, R.M., and Kaplan, E. (1991). Fine structure of receptive-field centers of X and Y cells of the cat. Vis. Neurosci. 6, 621–628.

Shah, N., Brackbill, N., Rhoades, C., Tikidji-Hamburyan, A., Goetz, G., Litke, A., Sher, A., Simoncelli, E., Chichilnisky, E.J. (2018). Inference of Nonlinear Spatial Subunits by Spike-Triggered Clustering in Primate Retina. bioRxiv. http://dx.doi.org/10.1101/496422.

Tien, N.-W., Pearson, J.T., Heller, C.R., Demas, J., and Kerschensteiner, D. (2015). Genetically Identified Suppressed-by-Contrast Retinal Ganglion Cells Reliably Signal Self-Generated Visual Stimuli. J. Neurosci. 35, 10815–10820.

Tsukamoto, Y., and Omi, N. (2015). OFF bipolar cells in macaque retina: type-specific connectivity in the outer and inner synaptic layers. Front. Neuroanat. 9, 122.

Tsukamoto, Y., and Omi, N. (2016). ON Bipolar Cells in Macaque Retina: Type-Specific Synaptic Connectivity with Special Reference to OFF Counterparts. Front. Neuroanat. 10, 104.

Victor, J.D., and Shapley, R.M. (1979). Receptive field mechanisms of cat X and Y retinal ganglion cells. J. Gen. Physiol. 74, 275–298.

Wässle, H. (2004). Parallel processing in the mammalian retina. Nat. Rev. Neurosci. 5, 747–757.

Wassle, H., Peichl, L., and Boycott, B.B. (1981). Morphology and Topography of on- and off-Alpha Cells in the Cat Retina. Proceedings of the Royal Society B: Biological Sciences 212, 157–175.

Watanabe, M., and Rodieck, R.W. (1989). Parasol and midget ganglion cells of the primate retina. J. Comp. Neurol. 289, 434–454.

Yamada, E.S., Bordt, A.S., and Marshak, D.W. (2005). Wide-field ganglion cells in macaque retinas. Vis. Neurosci. 22, 383–393.

Zelnik-Manor, L. and Perona, P. (2004). Self-Tuning Spectral Clustering. 17th Conference of Neural Information Processing Systems.

